# Genetic factors influence behavioural repeatability in juvenile poison frogs

**DOI:** 10.1101/2022.09.19.508512

**Authors:** Ria Sonnleitner, Emmi Alanen, Chloe Fouilloux, Janne K. Valkonen, Bibiana Rojas

## Abstract

Individual behaviour is a combination of previous experiences and genetic factors whose interaction can be adaptively adjusted to respond to changes in the surrounding environment. Understanding the continuity of behaviours both within and among individuals can help us disentangle the ecological and evolutionary significance underlying patterns of aggression, activity, boldness, and cooperation. In this study, we examined whether there is repeatability in the activity levels of juvenile dyeing poison frogs (*Dendrobates tinctorius*). This tropical species, known for its striking coloration and skin toxins, changes upon metamorphosis from an aquatic to a terrestrial habitat where individuals must immediately become active foragers to acquire their chemical defences. We did not find individual behaviour to be repeatable, however, we detected repeatability in activity at the family level, suggesting that behavioural variation may be explained, at least partially, by genetic factors in addition to a common environment. The assumption that activity level is very important for the survival of individuals after metamorphosis is supported by our results in that genetic factors seem to play a role in the formation of activity level. Further experiments are needed to investigate to what extent other areas of the individuals’ lives are affected by the respective activity levels, and what significance this has for the survival of a population.

## Introduction

Behaviour is shaped by internal (genetics, physiology; Robinson et al., 2008) and external (environment, previous experiences; Beach & Jaynes, 1954; Salvanes et al., 2007) factors. Because no two individuals are exposed to exactly the same environmental and physiological conditions, there is inter-individual variation in behaviour. From an evolutionary perspective, different behaviours may be targets of selection (Dall et al., 2004; Shelton & Martins, 2017); for example, in three-spined sticklebacks (*Gasterosteus aculeatus*), bold individuals outcompete shy individuals in catching prey (Ward et al., 2004), which may be critical under food-limited conditions, whereas in common lizards (*Lacerta vivipara*), the more active individuals appear to be less susceptible to parasites (Clobert et al., 2000), which may significantly affect their fitness. While behavioural variation within a species or a population is commonplace, individuals often behave the same way repeatedly, both across time and in familiar environments. Here, we define this repeatability as ‘behavioural consistency’, where individual behaviour is consistent across contexts and/or across different phases of life history (Schuster et al., 2017). From a statistical point of view, in contrast, ‘repeatability’ is defined as the proportion of variance between individuals from observations within the same context and within the same life history stage relative to the total variation in the trait in a population (Falconer, 1960; Lessells & Boag, 1987; Boake, 1989; Kamel & Mrosovsky, 2005; Schuster et al., 2017). Therefore, behaviours that have relatively low intra-individual variance compared to high inter-individual variance can be considered to be more repeatable (Bell et al., 2009).

Repeatability has been shown to occur in a wide variety of taxa and in multiple functional classes of behaviour such as activity (amphibians: Smith & Doupnik, 2005; birds: Kralj-Fišer et al., 2007; insects: Nemiroff & Despland, 2007), aggression (birds: Garamszegi et al., 2006; Kralj-Fišer et al., 2007; fishes: Bakker, 1986; Riddell & Swain, 1991; insects: Clark & Moore, 1995; Brown et al., 2006), courtship (amphibians: Ryan & Rand, 2003; Smith & Hunter, 2005; birds: Forstmeier & Birkhead, 2004; spiders: Rivero et al., 2000), foraging (fishes: Martins et al., 2005; insects: Missoweit et al., 2007; mammals: Koteja et al., 2003) and habitat selection (reptiles: Kamel & Mrosovsky, 2004 & 2005). However, in a meta-analysis of a large number of published studies on behavioural repeatability, Bell et al. (2009) found that not all behaviours are equally repeatable: behaviours constrained by animal physiology and morphology are more repeatable than behaviours that are sensitive to an animal’s social environment or energetic needs (Castellano et al., 2002; Smith and Hunter, 2005; Bell et al., 2009). In addition, individual experience may also play a particular role in the emergence of behavioural repeatability (Urszán et al., 2015).

Amphibians provide an excellent opportunity for studies on behavioural repeatability as metamorphosis entails extreme changes in habitat and ecological niche. Consequently, research on amphibians can provide important insights into how behaviours are related to and may arise from both occupying diverse ecologies as well as changes in physiology and morphology throughout development (Wilson & Krause, 2012). Although amphibians are known for their plasticity, especially during their larval stages (Wilbur & Collins, 1973; Smith-Gill & Berven, 1979; Zakrzewski, 1990; Nicieza, 2000; Buskirk, 2009; Cuello et al., 2014), there are numerous studies revealing behavioural repeatability in larvae (Smith & Doupnik, 2005; Wilson & Krause, 2012; Brodin et al., 2013), juveniles (Koenig & Ousterhout, 2018), and adults (Ryan & Rand, 2003; Smith & Hunter, 2005), in traits such as boldness (*Rana ridibunda*: Wilson & Krause, 2012; *Rana temporaria*: Brodin et al., 2013), exploration (*Rana ridibunda*: Wilson & Krause, 2012; *Rana catesbeiana*: Carlson & Langkilde, 2013), courtship (*Physalaemus pustulosus*: Ryan & Rand, 2003; *Litoria booroolongensis*: Smith & Hunter, 2005) and activity (*Rana catesbeiana*: Smith & Doupnik, 2005; *Rana ridibunda*: Wilson & Krause, 2012; *Ambystoma maculatum*: Koenig & Ousterhout, 2018).

While many studies have long addressed the diverse and complex behavioural characteristics of poison frogs, only recent studies have addressed the repeatability of their behaviours. For example, Chaloupka et al. (2022) found moderate repeatability in the territorial aggressiveness of the brilliant-thighed poison frog, *Allobates femoralis*. Here, we investigated the repeatability of activity in recently metamorphosed individuals of the dyeing poison frog, *Dendrobates tinctorius*. This species is well known for its warning coloration (Wollenberg et al., 2008; Rojas & Endler, 2013) and the possession of skin toxins (Saporito et al., 2012; Lawrence et al., 2019) sequestered from their specialised diet consisting mainly of ants and mites (Saporito et al., 2007, 2012). Together, these characteristics provide them with great protection from predators (via aposematism; Poulton, 1890) from the end of their aquatic life phase: upon metamorphosis, individuals go through not only dramatic morphological changes, but also a change in habitat. In the process, individuals must emerge from their pools to become active foragers from one day to the next (Toft, 1995; Caldwell, 1996; Pacheco et al., 2021). It has been shown that certain behavioural traits such as boldness and activity can predict the dispersal and survival of individuals after reintroduction into the wild (Hammond et al., 2021), and it is very likely that this is also applicable for the time after metamorphosis. Activity is vital for froglets not only to avoid starvation, but also to acquire the nutritional sources on which they are dependent to sequester precursors of their chemical defences (i.e., skin alkaloids; Saporito et al., 2007, 2012). However, increased activity may also increase detection risk by predators, who have demonstrated a preference for small frogs (Flores et al., 2015). Thus, the survival of young frogs is a balance between the benefits of active foraging and the cost of increased predation risk. Because the activity optima in young frogs is narrow, where falling below the ideal threshold might lead to starvation and above the ideal threshold might lead to increased predation risk, we expect a genetic component to individual activity to be detected through the repeatability of behaviour. Here, we used a genetically-controlled design to explore whether activity was repeatable at an individual and/or family level. We predicted that individual frogs would exhibit behavioural repeatability in two successive trials, and that closely related individuals would be similar in their activity levels, but different from unrelated individuals. This should give us insight into the extent to which behavioural strategies, especially survival strategies of metamorphs after leaving their old habitat and entering a new one, are repeatedly used by an individual and may be partly genetically determined.

## Methods

### Study species and husbandry

*Dendrobates tinctorius* is a poison frog species (family Dendrobatidae) native to tropical rainforests in the lowlands of the Guiana Shield. They are aposematic (i.e., display warning coloration and possess skin toxins; Noonan & Comeault 2009; Rojas et al. 2014a; Lawrence et al. 2019) and males exhibit an elaborate parental care behaviour. Recently-hatched tadpoles are transported on their father’s back from terrestrial clutches to water-filled plant structures at variable heights (Rojas 2014, 2015; Rojas & Pašukonis, 2019; Fouilloux et al. 2021). The tadpoles spend about two months unattended in these nurseries until metamorphosis, when they must start actively foraging in their new habitat on land (Rojas & Pašukonis, 2019).

The froglets used in this study were offspring from 12 breeding pairs kept in the laboratory of the University of Jyväskylä, Finland. The adult breeding pairs were housed in terrariums (115 L) equipped with expanded clay, leaf litter, moss substrate and hiding places at 26 °C (± 2 °C) and a humidity of 95 % and fed ad libitum with live, vitamin-coated fruit flies five times a week.

### Activity measurement

To determine the repeatability of activity in juvenile frogs, we measured latency to emerge, activity, resting and hiding (Table 1) of individuals across two trials. The time interval between the two trials was 13 - 14 days to avoid the risk of temporal autocorrelation (Koenig & Ousterhout, 2018). Activity was measured in an experimental arena which consisted of a transparent plastic box (size: 39 × 34 cm; height of walls: 9 cm) filled with 2 cm of soil. In the opposite corners of the arena were two shelters (size: 8 × 8 cm; height: 2 cm) whose positioning was randomised for each trial. At the beginning of each trial, individuals (N = 17) were weighed (weight: first experiment: X ± SD = 0.39 ± 0.06 g; second experiment: X ± SD = 0.39 ± 0.08 g), and then placed in a dark film canister (diameter: 3 cm; height: 2 cm) in the centre of the experimental arena, where they were kept for a 15-minute acclimation period. Following acclimation, the lid of the canister was removed and the frog was able to move around the arena freely. We recorded the time it took the frog to leave the box and observed the behaviour of the individual (latency to emerge/activity (= moving + climbing)/resting/hiding, Table 2) for 30 minutes, at 15-second intervals.

**Table 1.** Ethogram of juvenile behaviours.

| Behaviour | Definition |
| --- | --- |
| latency to emerge | The time it took the individual to leave the film canister at the beginning of the trial |
| activity | The sum of moving and climbing behaviours |
| resting | Individual is in one place; includes small changes of body position |
| hiding | Individual is positioned in a hiding place |

**Table 2.** Summary of the adjusted repeatability estimates (*R*, link scale) of froglet behaviour.

| *Variable* | *Level* | *R* | *SE* | *CI* | *p* |
| --- | --- | --- | --- | --- | --- |
| **Activity**<br>(moving + climbing) | Fixed | 0.190 | 0.211 | [0.043, 0.911] | - |
|  | Individual | 0 | 0.12 | [0, 0.417] | 1 |
|  | **Family** | **0.486** | **0.211** | **[0, 0.737]** | **0.001** |
| **Resting**<br>(in exposed areas) | Fixed | 0.157 | 0.165 | [0.032, 0.650] | - |
|  | **Individual** | **0.402** | **0.214** | **[0, 0.731]** | **0.045** |
|  | Family | 0 | 0.088 | [0, 0.310] | 0.5 |
| **Latency to emerge** | Fixed | 0.005 | 0.0684 | [0.001, 0.259] | - |
|  | Individual | 0.0239 | 0.136 | [0, 0.463] | 0.470 |
|  | Family | 0.387 | 0.197 | [0, 0.694] | 0.093 |

Eighteen juveniles from six different families were tested with three individuals per family. One froglet was excluded from analysis due to abnormal physical condition observed throughout its second trial. Thus, repeatability data were available for 17 juveniles.

### Statistical analysis

In this study, we investigated whether the behaviour of individuals at the juvenile stage is repeatable between two trials, either at the individual or at the family level. Activity (the sum of moving and climbing counts), resting and hiding was coded as counts from 15-second scan intervals. Latency to emerge was coded as the time (seconds) an individual spent in the start box. Individual froglet identities nested within froglet families were taken as random effects across all models, which made models suitable for estimating variance components and calculating the repeatability of behaviour at individual and family levels. We used generalised linear mixed models (GLMM) to test the effects of trial number and weight on behaviour.

Models were built using the template model builder “glmmTMB” (Magnusson et al., 2020), and residual diagnostics and overdispersion were tested using the package “DHARMa” (Hartig, 2020). None of the final models was overdispersed or zero-inflated. Multiple families were fitted to each model, using AICc (package “AICcmodavg” (Mazerolle, 2020)) model family parameterization were ranked. Models with the lowest log-likelihood were chosen. GLMMs were fit with a negative binomial distribution with either a linear or quadratic parameterization.

The selected models were used for the behavioural repeatability analyses. Repeatability was estimated using the package “rptR” (Stoffel et al., 2019), where within- and between-group variance was captured by intra-class correlation (ICC) coefficients. Here, we were interested in both the variance explained by the random effects (individual and family) as well as model fixed effects (weight, trial). We therefore used *adjusted repeatabilities*, where repeatability is measured after controlling for fixed effects (Nakagawa & Schielzeth, 2010). The hiding (shelter) variable was fully separated at two family levels (where families 11 and 15 never hid in the shelters) and strongly correlated with other response variables making it unreliable for repeatability measures. Uncertainty of the adjusted repeatability estimates (CI) came from 1000 parametric bootstraps and 1000 permutations. Repeatability estimates of count variables were reported as link scale approximations because they were fit using a log link. All analyses were conducted with R version 4.1.1 (R Core Team, 2020).

## Results

With respect to our general models investigating the effects of froglet mass and trial number on behaviour, we found that overall froglet activity significantly decreased across trials (GLMM, *Z* = −2.47, p = 0.014, Supp. Table 3), but no overall trial effect on hiding (GLMM, *Z* = 1.49, p = 0.134) or latency to emerge (GLMM, z-val = −0.540, p = 0.588).

Hiding models revealed high between-subject variance on the family level (τ_00_ = 11.23), which indicates a notable measure in how much families differed from each other. This is generated by complete separation of the variable across two froglet families (Fam. 11 and 15, see Supp. Fig 3) that never exhibited hiding behaviour across either trials. Across all models froglet weight did not significantly impact any behaviour.

When exploring the repeatability of froglets through both individual and family-level groupings, we find that froglet activity was repeatable at the family level (R = 0.486, p < 0.001, Fig. 1). Surprisingly, we found that on an individual level resting behaviour was repeatable (R = 0.402, p = 0.045); we did not detect any repeatability on either individual or family levels with respect to latency to emerge from the starting shelter.

**Figure 1.**
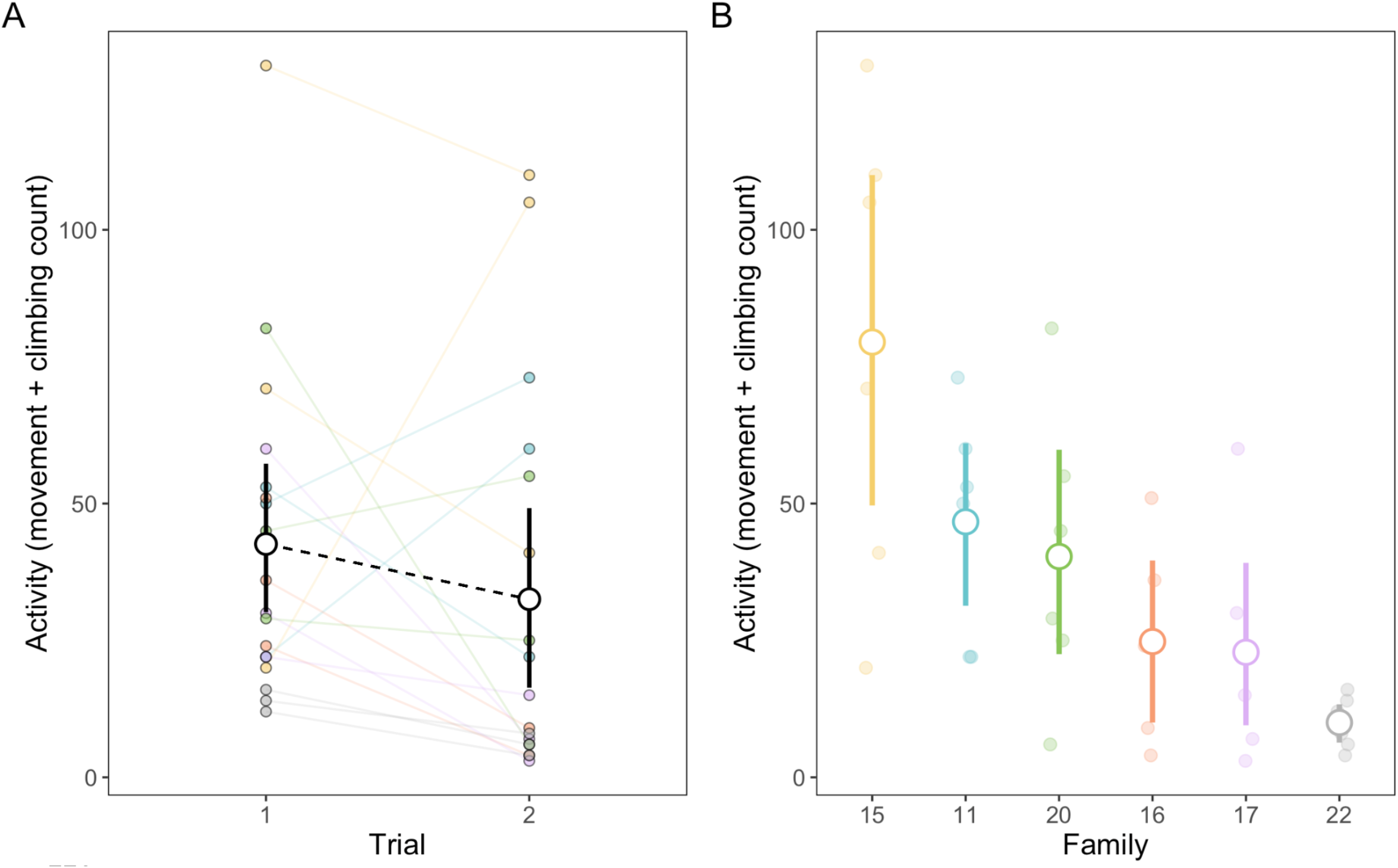
Activity of juvenile frogs across two trials. (A) Overall activity decreased across trials, where juveniles were more active in the first trial than in the second. Connected colored points represent individual activity in each trial and black pointranges represent trial averages. (B) Activity was repeatable across families, where some families were repeatedly more active than others (R = 0.473, p = 0.001). Underlying coloured points represent individual activity (grouped by family) in each trial. Point ranges illustrate means and 95% C.I.

**Figure 2.**
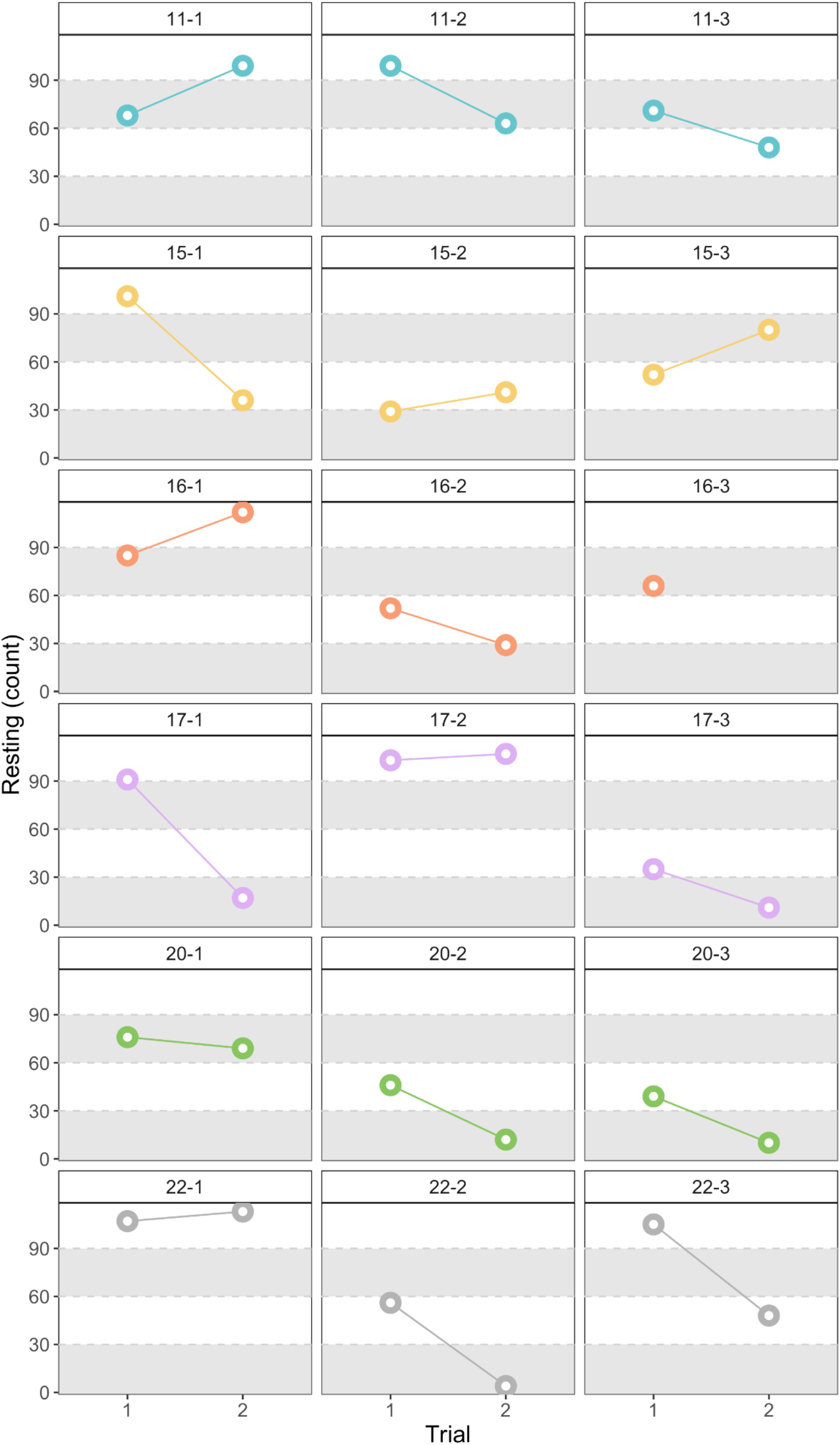
Repeatability of resting behaviour. Froglets demonstrate repeatable levels of resting behaviour on an individual level. Each panel represents an individual, each row consists of a family; coloured lines connect count between both trials to facilitate interpretation. Note individual 16-3 was only measured once but was included for the calculation of family-level effects.

## Discussion

Activity is an important component of an individual’s behaviour which provides a meaningful correlate for predicting survival in the wild (Hammond et al., 2021). In poison frogs, activity is important for young froglets, as they must become active foragers in their new terrestrial habitat right after metamorphosis, both for immediate sustenance and to sequester the alkaloids necessary for their chemical defence. Here, we explore the individual activity patterns in *Dendrobates tinctorius* froglets and test to what degree these behaviours are repeatable.

The repeatability of activity is not uncommon across animal taxa and has been identified in birds (e.g., Kralj-Fišer et al., 2007; Kluen & Brommer, 2013; Wuerz & Krüger, 2015), insects (e.g., Nemiroff & Despland, 2007; Herde & Eccard, 2013; Wexler et al., 2016; Videlier et al., 2019), mammals (e.g., Twiss & Franklin, 2010; Menzies et al., 2013; Schuster et al., 2017; Santicchia et al., 2021), fish (e.g., Poulin et al., 2012; Fürtbauer et al., 2015; White et al., 2015; Thomson et al., 2020), and frogs (e.g., Smith & Doupnik, 2005; Wilson & Krause, 2012; Koenig & Ousterhout, 2018). Individual activity is closely linked to foraging success in ant-preying poison frogs (Toft, 1981), and successful foraging is imperative for both short-term survival (Marcus, 1981; Donoghue, 1998) and long-term protection via toxin accumulation (Santos et al., 2003; Darst et al., 2005). Thus, we hypothesised that activity would be repeatable on an individual level, where some individuals would consistently be more active than others. Likewise, we predicted that the variation in individual behaviour would be more likely to be captured by the repeatability of behaviour within families, with similar mean levels of activity observed between close relatives that could be due to a shared genetic component. Contrary to our hypothesis, we found no repeatability at the individual level. We did, however, find repeatability at the family level, where some families were consistently more active than others. Our results show that the behaviour of a group of relatives is repeatable, even though the behaviour of an individual may not be, suggesting that some of the behavioural variation may be explained by genetic factors when individuals are reared under similar conditions. Family identity (i.e., genetic factors, maternal effects) has been shown to affect various aspects of life, such as growth rate and activity in wood frogs (Warne et al., 2013; Relyea, 2005), resting metabolic rate in alpine newts (Baškiera et al., 2021), call variation in big brown bats (Masters et al., 1995), feeding behaviour in red-backed salamanders (Gibbons et al. 2005), and escape-related behaviour in voles (Lantová et al. 2011).

From an evolutionary perspective, it seems logical that multiple activity strategies exist and are maintained within populations, as different foraging strategies may be preferred depending on local predator and prey density. Thereby families with increased activity are likely able to find food more efficiently at the potential cost of increasing detectability by predators. In contrast, individuals who are less active and therefore might take longer to consume a given amount of food would lack chemical protection for a longer time period, but at the same time are less likely to be detected by predators due to reduced activity. Therefore, we assume that when prey density and/or the number of predators are low, the more active individuals will have advantages, whereas when prey density and/or the number of local predators are high, less active individuals will benefit, thus, successfully maintaining both strategies within a population. For future studies, it will be interesting to investigate whether the respective strategy regarding activity levels and foraging style is also influenced by the individual colour pattern, as it has already been established that the dorsal colour patterns of *D. tinctorius* may provide protection against predators (Noonan & Comeault, 2009; Comeault & Noonan, 2011; Rojas et al. 2014a; Lawrence et al 2019), and that there is a relationship between dorsal colour pattern and movement type (fast and straight vs. slow and random; Rojas et al., 2014b).

With respect to the repeatability of activity, we found that, overall, the activity of froglets decreased significantly across two trials; no repeated measurement effects were detected in other measures of behaviour such as hiding or latency to emerge. The decreased activity in the second trial could be related to the recognition of the experimental environment after prior exposure during the first run. Chajma et al. (2020) similarly found reduced activity in smooth newts on all repeat tests as a possible consequence of the animals’ memory of the arena.

In some amphibians, the expression of certain behaviours has been shown to remain consistent from the tadpole stage through metamorphosis and into adulthood (i.e., consistency; Wilson & Krause, 2012; Koenig & Ousterhout, 2018). For future studies, it would be interesting to examine the consistency of different behaviours throughout the life stages of *D. tinctorius* to gain deeper insight into the stability of behavioural syndromes in the face of profound physiological, morphological, and ecological changes implicit in animals with complex life cycles (Wilson and Krause, 2012).

## Conclusion

We conclude that there are different strategies in *D. tinctorius* populations regarding the level of activity when exploring a new habitat. These activity levels might play a major role in their survival rates and seem to be influenced by genetic factors. Further experiments are needed to investigate to what extent other areas of the individuals’ lives are affected by the respective activity levels, and what significance this has for the survival of a population.

## Supplementary Materials

**Supp Table 3.**
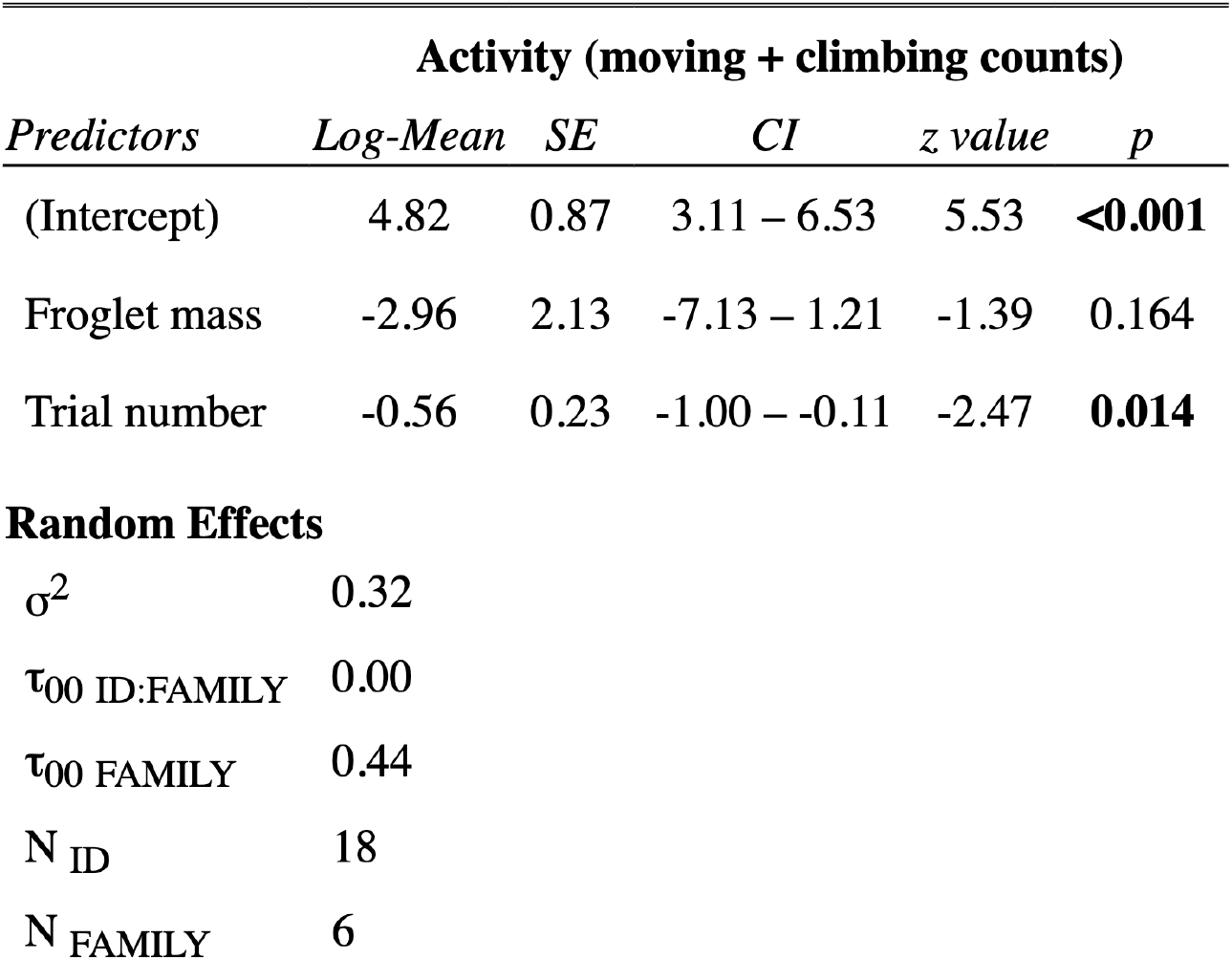
GLMM model output of froglet activity. Model included nested random effect of Family/Froglet ID, and fitted with a negative binomial family with a quadratic parameterization.

**Supp Table 4.** GLMM model output of froglet hiding. Model included nested random effect of Family/Froglet ID, and fitted with a negative binomial family with a quadratic parameterization. Note high random between-family variance (τ_00_ = 11.23).

| Hiding (counts) |  |  |  |  |  |
| --- | --- | --- | --- | --- | --- |
| *Predictors* | *Log-Mean* | *SE* | *CI* | *z value* | *p* |
| (Intercept) | -4.78 | 5.05 | -14.69 – 5.12 | -0.95 | 0.344 |
| Froglet mass | 13.80 | 11.52 | -8.78 – 36.38 | 1.20 | 0.231 |
| Trial number | 1.49 | 0.99 | -0.46 – 3.43 | 1.50 | 0.134 |
| **Random Effects** |  |  |  |  |  |
| $\sigma^2$ | 1.89 | | | | |
| $\tau_{00}$ ID:FAMILY | 0.00 | | | | |
| $\tau_{00}$ FAMILY | 11.23 | | | | |
| N ID | 18 |  |  |  |  |
| N FAMILY | 6 |  |  |  |  |

**Supp. Fig 3.**
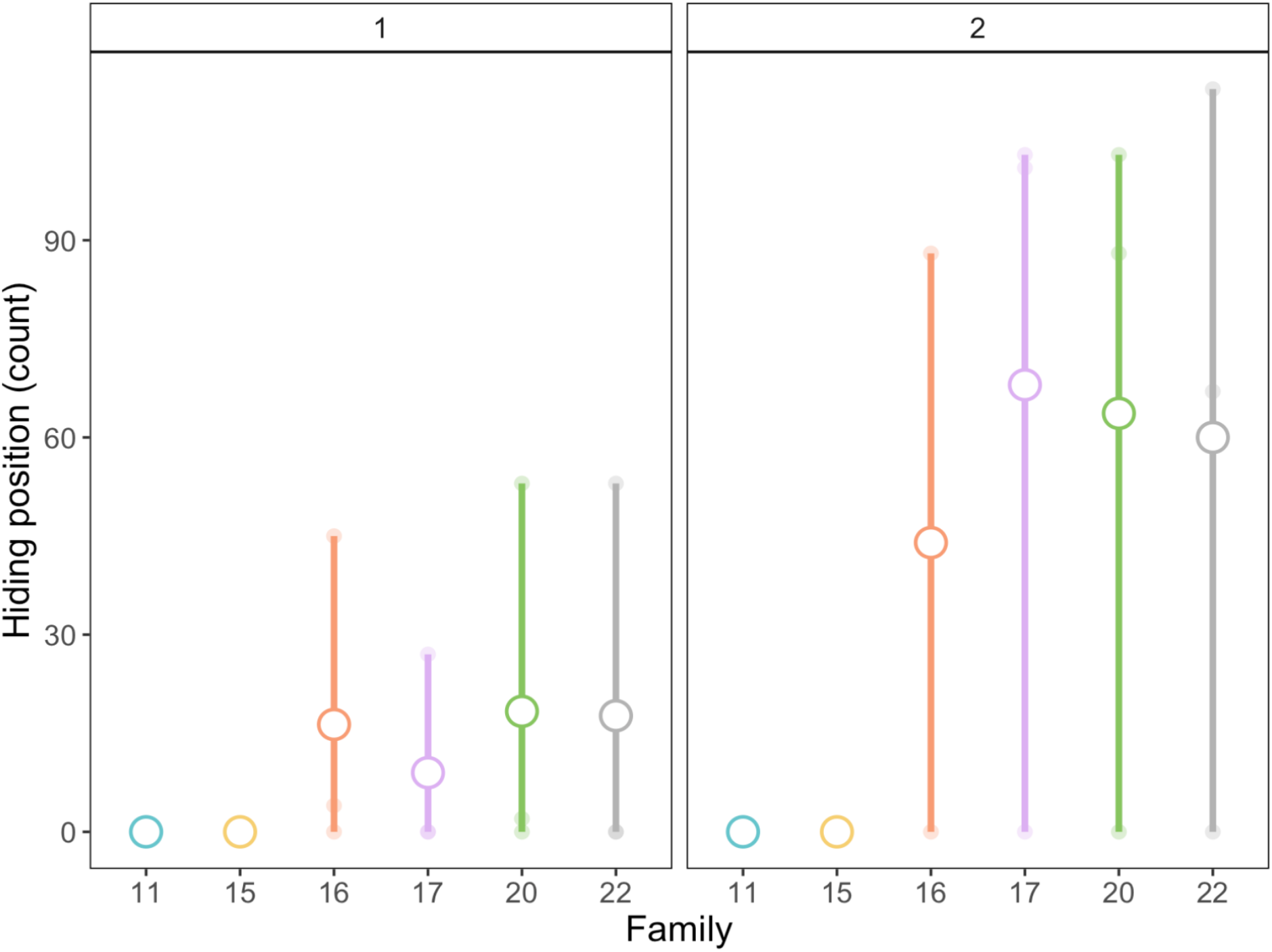
Hiding behaviour by family. Note the between-group variance of behaviour on the family level.

**Supp Table 5.**
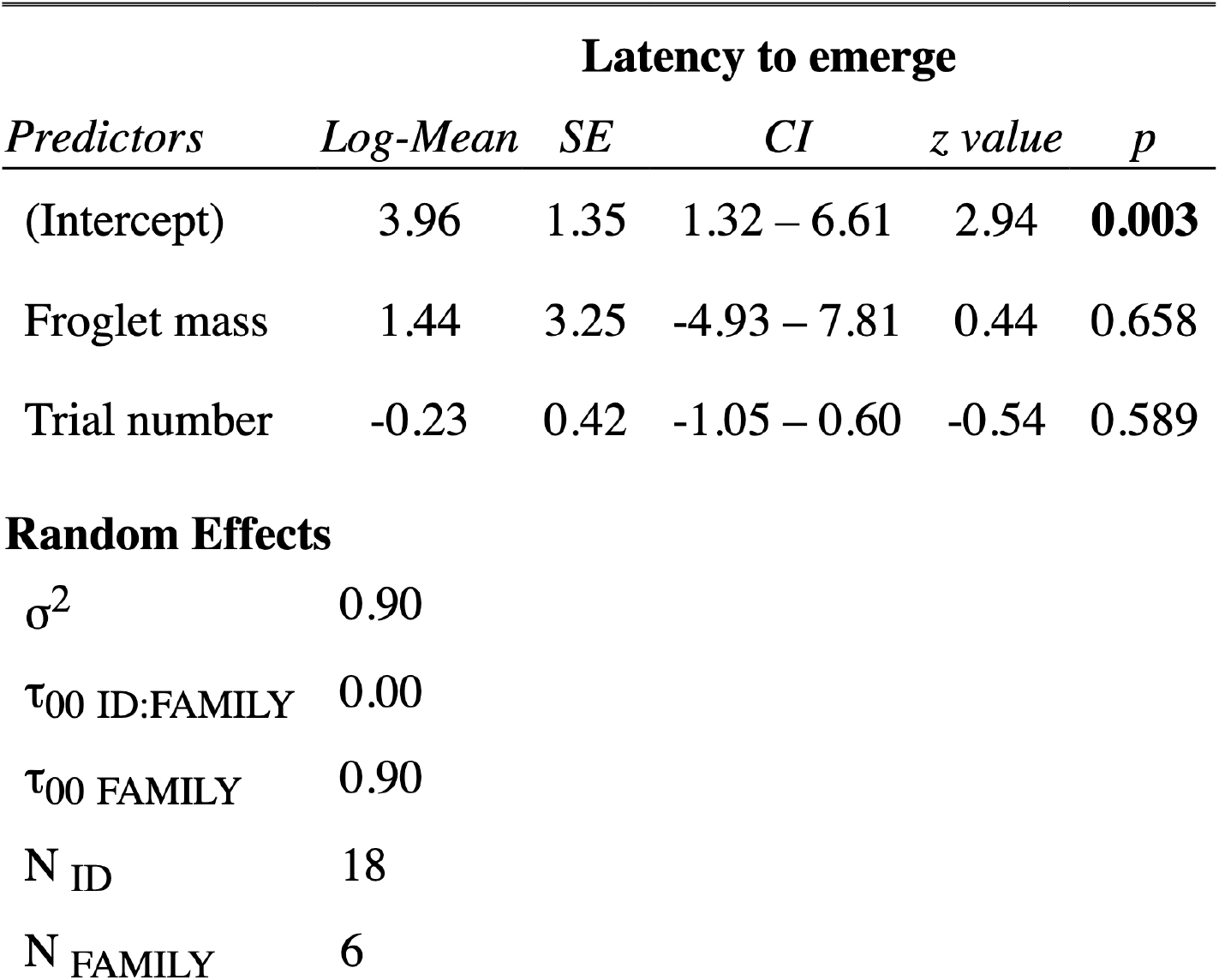
GLMM model output of froglet latency to emerge. Model included nested random effect of Family/Froglet ID, and fitted with a negative binomial family with a quadratic parameterization.

| Latency to emerge |  |  |  |  |  |
| --- | --- | --- | --- | --- | --- |
| *Predictors* | *Log-Mean* | *SE* | *CI* | *z value* | *p* |
| (Intercept) | 3.96 | 1.35 | 1.32 – 6.61 | 2.94 | **0.003** |
| Froglet mass | 1.44 | 3.25 | -4.93 – 7.81 | 0.44 | 0.658 |
| Trial number | -0.23 | 0.42 | -1.05 – 0.60 | -0.54 | 0.589 |
| **Random Effects** |  |  |  |  |  |
| $\sigma^2$ | 0.90 | | | | |
| $\tau_{00}$ ID:FAMILY | 0.00 | | | | |
| $\tau_{00}$ FAMILY | 0.90 | | | | |
| N ID | 18 |  |  |  |  |
| N FAMILY | 6 |  |  |  |  |

## Author contributions

**Ria Sonnleitnner:** Data curation (supporting); Writing-original draft (lead). **Emmi Alanen:** Investigation (lead); Data curation (lead); Formal analysis (supporting); Writing-review & editing (supporting). **Chloe Fouilloux:** Conceptualization (equal); Methodology (equal); Investigation (supporting); Formal analysis (lead); Supervision (equal); Writing-review & editing (equal). **Janne K. Valkonen:** Conceptualization (equal); Methodology (equal); Investigation (supporting); Formal analysis (supporting); Supervision (equal); **Bibiana Rojas:** Conceptualization (equal); Methodology (supporting); Funding acquisition (lead); Supervision (equal); Writing-review & editing (equal).

## Ethics

This study was approved by the National Animal Experiment Board at the Regional State Administrative Agency for Southern Finland (ESAVI/9114/04.10.07/2014) and followed the Association for the Study of Animal Behaviour’s guidelines for the treatment of animals in behavioural research and teaching (ASAB, 2018).

## Data accessibility

A data file can be found in the electronic supplementary materials.

## Acknowledgments

We are grateful to Teemu Tuomaala for help with frog care, and to Carita Lindstedt Silva Uusi-Heikkilä, and Carolin Dittrich for thoughtful feedback on earlier versions of the manuscript. We highly value equity, diversity, and inclusion in science, and therefore our study was done taking into account EDI best practice. We cherish the international and diverse nature of our team, which includes researchers from (5) different countries (Austria, Finland, France, United States and Colombia), backgrounds, and career stages, as it significantly contributed to the fulfilment and quality of the present study. This study was funded by the Academy of Finland (Academy Research Fellowship No. 319949 to BR).

